# Expansion of SARS-CoV-2-specific Antibody-secreting Cells and Generation of Neutralizing Antibodies in Hospitalized COVID-19 Patients

**DOI:** 10.1101/2020.05.28.118729

**Authors:** Renata Varnaitė, Marina García, Hedvig Glans, Kimia T. Maleki, John Tyler Sandberg, Janne Tynell, Wanda Christ, Nina Lagerqvist, Hilmir Asgeirsson, Hans-Gustaf Ljunggren, Gustaf Ahlén, Lars Frelin, Matti Sällberg, Kim Blom, Jonas Klingström, Sara Gredmark-Russ

**Author notes:** H.G., K.T.M. and J.T.S. contributed equally to this work. **Corresponding author** Sara Gredmark-Russ, Center for Infectious Medicine, ANA Futura, Department of Medicine, Karolinska Institutet, Karolinska University Hospital Huddinge, 141 52 Stockholm, Sweden.

## Abstract

Coronavirus disease 2019 (COVID-19), caused by severe acute respiratory syndrome coronavirus 2 (SARS-CoV-2), emerged in late 2019 and has since become a global pandemic. Pathogen-specific antibodies are typically a major predictor of protective immunity, yet B cell and antibody responses during COVID-19 are not fully understood. Here, we analyzed antibody-secreting cell (ASC) and antibody responses in twenty hospitalized COVID-19 patients. The patients exhibited typical symptoms of COVID-19, and presented with reduced lymphocyte numbers and increased T cell and B cell activation. Importantly, we detected an expansion of SARS-CoV-2 nucleocapsid protein-specific ASCs in all twenty COVID-19 patients using a multicolor FluoroSpot assay. Out of the 20 patients, 16 had developed SARS-CoV-2-neutralizing antibodies by the time of inclusion in the study. SARS-CoV-2-specific IgA, IgG and IgM antibody levels positively correlated with SARS-CoV-2-neutralizing antibody titers, suggesting that SARS-CoV-2-specific antibody levels may reflect the titers of neutralizing antibodies in COVID-19 patients during the acute phase of infection. Lastly, we showed that interleukin 6 (IL-6) and C-reactive protein (CRP) concentrations were higher in serum of patients who were hospitalized for longer, supporting the recent observations that IL-6 and CRP could be used to predict COVID-19 severity. Altogether, this study constitutes a detailed description of clinical and immunological parameters in twenty COVID-19 patients, with a focus on B cell and antibody responses, and provides tools to study immune responses to SARS-CoV-2 infection and vaccination.

## Introduction

Coronavirus disease 2019 (COVID-19), caused by severe acute respiratory syndrome coronavirus 2 (SARS-CoV-2), emerged in late 2019 and has since become a global pandemic (1). SARS-CoV-2 primarily affects the respiratory tract and can cause severe respiratory illness, which sometimes requires mechanical ventilation (2). The world is racing to understand the immune responses to SARS-CoV-2 in order to identify correlates of protection, and to aid treatment and vaccine development. In the majority of infections, pathogen-specific antibodies are one of the main contributors to protective immunity, yet B cell and antibody responses during COVID-19 are currently not fully understood.

B cells are one of the key players of the adaptive immune system and can provide sterilizing immunity against pathogens through the secretion of pathogen-specific antibodies. B cells carry immunoglobulin (Ig) surface receptors with a high range of specificity that can directly recognize different antigens. Early during acute infections upon the encounter with the antigen, B cells differentiate into antibody-secreting cells (ASCs), including plasmablasts and plasma cells, which produce large quantities of pathogen-specific antibodies (3).

ASCs can expand to very high levels during some acute human infections (4-7). For example, during human dengue virus infection, ASCs expand to constitute an average of 47% of all circulating B cells (5). Meanwhile, at steady state conditions in healthy individuals, ASCs comprise only up to around 1% of all circulating B cells (5). The ASC response is typically transient, peaking at around day 7 of the febrile stage of infection (8). However, during acute human respiratory syncytial virus (RSV) infection, ASC response could be detected for as long as 22-45 days after the onset of symptoms, which was found to be associated with prolonged shedding of RSV in the airways (9). Thus, the kinetics of ASC response may be dependent on pathogen persistence.

In humans, ASCs can be identified using flow cytometry as CD19^+^CD20^low/–^ isotype-switched B cells with high cell surface expression of CD38 and CD27 molecules, as well as intracellular expression of proliferation marker Ki-67 (10, 11). Although flow cytometry is a valuable tool to measure the frequencies of total ASCs in peripheral blood, it only allows phenotypic characterization. Functional antibody secretion capacity and antigen-specificity of ASCs are typically described using the gold standard enzyme-linked immunospot (ELISpot) assay (12). ELISpot is a highly sensitive method allowing for detection of ASCs producing different antibody isotypes at single-cell level, including IgA, IgG and IgM, against different antigens. Flow cytometry and ELISpot can therefore be used in combination to comprehensively describe the magnitude and specificity of B cell responses to infectious agents.

Antigen-specific antibodies can contribute to pathogen control either by (i) directly interfering with the entry of a pathogen into the target cells (referred to as neutralization); or by (ii) assisting the effector cells in recognizing and eliminating infected target cells (known as antibody-dependent cellular cytotoxicity) (13). Neutralizing antibodies are typically considered to be the strongest predictor of protective immunity following natural infection or vaccination. Therefore, it is of great importance to evaluate the neutralizing capacity and persistence of specific antibodies in acute viral infections such as SARS-CoV-2.

Here, we provide a detailed description of early B cell and antibody responses to SARS-CoV-2 infection in a cohort of twenty hospitalized patients with severe COVID-19. We found a substantial level of ASCs expansion in all patients, and that not all patients developed detectable levels of SARS-CoV-2-specific and neutralizing antibodies at the time of sampling. We also show increased levels of adaptive immune system activation and increased serum concentrations of inflammatory markers interleukin 6 (IL-6) and C-reactive protein (CRP), the latter two correlating with the duration of hospitalization in the present COVID-19 patient cohort. This study constitutes a detailed description of clinical and immunological B cell-related parameters in COVID-19 patients, and describes tools to study immune responses to SARS-CoV-2 infection and vaccination.

## Materials and Methods

### Ethics statement

The study was approved by the Regional Ethical Review Board in Stockholm, Sweden and by the Swedish Ethical Review Authority. All COVID-19 patients and healthy controls included in this study provided a written informed consent for participation.

### Study subjects and sampling of peripheral blood

Peripheral blood samples were collected from 20 adult COVID-19 patients hospitalized in April 2020 at the Karolinska University Hospital in Stockholm, Sweden (5 females and 15 males; age range between 34 and 67 years; median age 53 years) (Table I). Patients were diagnosed with COVID-19 by RT-PCR for SARS-CoV-2 in either nasopharyngeal swabs (18/20 patients) or sputum (2/20 patients). Diagnostics were performed at the diagnostic laboratory at the Karolinska University Hospital, Stockholm, Sweden. Peripheral blood samples from patients were taken at a median of 15 days after self-reported onset of symptoms (range 7-19 days). Peripheral blood samples of 7 healthy controls were collected in parallel (2 females and 5 males; age range between 26 and 53 years; median age 31 years).

**Table I.** Clinical characteristics of 20 hospitalized COVID-19 patients.

| **Cohort characteristics** |  |
| --- | --- |
| Age, years, median (range) | 53 (34-67) |
| Male, n (%) | 15 (75) |
| Symptom onset to sampling, days, median (range) | 15 (7-19) |
| Hospitalization to sampling, days, median (range) | 3 (2-10) |
| Symptom onset to discharge, days, median (range) | 19.5 (9-30) |
| Duration of hospitalization, days, median (range) | 7.5 (3-25) |
| Intensive care unit (ICU) treatment<sup>a</sup>, n (%) | 3 (15) |
| Fatal outcome, n (%) | 1 (5) |
| **Comorbidities** | **n (%)** |
| Overweight (BMI >25), n/N (%) | 14/14 (100)<sup>b</sup> |
| Diabetes mellitus type II | 4 (20) |
| Hypertension | 3 (15) |
| Asthma | 2 (10) |
| Cardiovascular disease | 1 (5) |
| **Symptoms** | **n (%)** |
| Fever (>38°C) | 20 (100) |
| Cough | 20 (100) |
| Respiratory difficulties (SpO<sub>2</sub> ≤ 93% on room air) | 20 (100) |
| Myalgia | 9 (45) |
| Chest pain | 6 (30) |
| Diarrhea | 4 (20) |
| **Treatment** | **n (%)** |
| Supplemental oxygen<sup>c</sup> | 20 (100) |
| High flow nasal oxygen | 4 (20) |
| Inhaled bronchodilators | 2 (10) |
| Low molecular weight heparin | 17 (85) |
| Antibiotics | 14 (70) |
| Immunomodulatory drugs<sup>d</sup> (given before sampling) | 3 (15) |

Abbreviations: n: number of patients; BMI: body mass index. SpO<sub>2</sub>: percutaneous oxygen saturation.
<sup>a</sup> ICU treatment before study sampling (n=2), ICU treatment after study sampling (n=1).
<sup>b</sup> n/N (%), where N is the total number of patients with available data.
<sup>c</sup> Oxygen flow rate – minimum 2.5 L/min. 1 patient in mechanical ventilation.
<sup>d</sup> Chloroquine phosphate (n=2) or Anakinra (n=1).

Peripheral blood mononuclear cells (PBMCs) were isolated from heparinized anti-coagulated blood using density gradient Lymphoprep medium (Stemcell Technologies) following the manufacturer’s instructions, and immediately used for flow cytometry and FluoroSpot assays. Serum was collected from COVID-19 patients and healthy controls in BD Vacutainer serum tubes with spray-coated silica (BD Biosciences). After coagulation for up to 2 hours at RT, serum was isolated by centrifugation at 2000 *g* for 10 min and immediately stored at −80 °C for later analysis.

### Absolute counts of lymphocytes in peripheral blood

Absolute numbers of CD45^+^, CD3^+^, CD4^+^, CD8^+^, and CD19^+^ cells in peripheral blood were measured using BD Trucount Tubes (BD Biosciences). 50 µL of anti-coagulated whole blood were added into Trucount Tubes within 3 hours after blood extraction and stained with either anti-CD45-PerCP (2D1), anti-CD3-FITC (SK7), anti-CD4-APC (SK3) and anti-CD8-PE (SK1), or anti-CD45-PerCP (2D1) and anti-CD19-AF488 (HIB19) (all from BioLegend). After 15 minutes of incubation at RT, stained whole blood was fixed and red blood cells lysed with 2X BD FACS Lysing Solution (BD Biosciences). Samples were acquired on a BD Accuri C6 Plus flow cytometer. Bead number recorded was used to quantify absolute CD45^+^, CD3^+^, CD4^+^, CD8^+^, and CD19^+^ cell counts per microliter of blood. ASC numbers per microliter of blood were calculated based on CD19^+^ B cell numbers measured by absolute cell counting and on frequencies of ASCs within CD19^+^ cells measured by flow cytometry.

### Flow cytometry

Staining with fluorescently-labelled antibodies was performed on freshly isolated PBMCs. Briefly, cells were incubated with surface staining antibodies diluted in PBS for 30 min at 4°C in the dark, followed by 3 washes with flow cytometry buffer (2% FCS and 2 mM EDTA in PBS). Cells were then fixed and permeabilized using eBioscience Foxp3/Transcription Factor Staining Buffer Set (Thermo Fisher Scientific) and later incubated with antibodies diluted in PBS for intracellular staining for 30 min at 4°C in the dark. Finally, samples were incubated in a 2% formaldehyde solution (Polysciences) for 2 h, washed and resuspended in flow cytometry buffer, and data subsequently acquired on a BD LSRFortessa flow cytometer equipped with 355, 405, 488, 561, and 639 nm lasers and BD FACSDiva Software (BD Biosciences). For a detailed gating strategy see Supplementary Fig. 1.

The following monoclonal antibody conjugates were used for cell surface staining: anti-CD8-Qdot605 (3B5) (Thermo Fisher Scientific), anti-CD19-BUV395 (SJ25C1), anti-CD14-V500 (MφP9), anti-CD4 -BUV737 (RPA-T4) (all from BD Biosciences), anti-CD123-BV510 (6H6), anti-CD27-BV650 (O323), anti-CD20-FITC (2H7), anti-CD38-BV421 (HB-7), anti-IgD-PE-Cy7 (IA6-2), anti-IgM-BV785 (MHM-88) (all from BioLegend), anti-CD3-PE-Cy5 (UCHT1), anti-CD56-ECD (N901) (all from Beckman Coulter), and anti-IgA-APC (REA1014) (Miltenyi). LIVE/DEAD Fixable Near-IR Dead Cell Stain Kit (Thermo Fisher Scientific) was used as a viability marker. The following monoclonal antibody conjugates were used for intracellular staining: anti-IgG-PE (HP6017) (BioLegend) and anti-Ki-67-AF700 (B56) (BD Biosciences).

### FluoroSpot assay for antibody-secreting cells

The number of SARS-CoV-2 nucleocapsid (N) protein-specific IgA, IgG and IgM antibody-secreting cells (ASCs), as well as the total number of IgA-, IgG- and IgM-ASCs in freshly isolated PBMCs were measured using a multicolor B cell FluoroSpot kit with modifications (Mabtech). Briefly, ethanol-activated IPFL membrane plates were coated overnight with either: (i) anti-IgG, anti-IgA, and anti-IgM capture antibodies (15µg/mL of each) for the detection of all ASCs, or (ii) SARS-CoV-2 N protein (10 µg/mL) for the detection of SARS-CoV-2-specific ASCs. The plates were washed with PBS and blocked with R10 media (RPMI-1640 with 10% FCS, 1% Pen/Strep, 2mM L-Glutamine (all from Thermo Fisher Scientific)) for 30 minutes at RT before the addition of freshly isolated PBMCs. Plates were then incubated at 37°C in 5% CO_2_ for 20 hours and then developed with anti-human IgG-550 (yellow fluorescence), anti-human IgA-490 (green fluorescence) and anti-human IgM-640 (red fluorescence) secondary detection antibodies (diluted 1:500 each) (all antibodies from Mabtech). Fluorescent spots indicating a single ASC were detected with an IRIS FluoroSpot reader and counted with Apex software (Mabtech).

### Recombinant SARS-CoV-2 nucleocapsid protein

A full-length nucleocapsid (N) phosphoprotein nucleotide sequence (1293 base-pairs) of the SARS-CoV-2 virus was optimized and synthesized (Genscript). The synthesized sequence was cloned into a PET-30a(+) vector with a carboxyterminal His tag for detection of protein expression in *E.coli*. The *E. coli* strain BL21 Star (DE3) was transformed with the recombinant plasmid and a single colony was inoculated into TB medium containing antibiotic and cultured at 37°C at 200 rpm and then induced with IPTG. Protein purity and molecular weight were determined by SDS-PAGE and Western blot according to standard procedures (Genscript).

### SARS-CoV-2 isolation from serum

50 µL of serum were mixed with 150 µL of EMEM (Gibco) and added to confluent Vero E6 cells seeded in 24-well plates. Cells were incubated with diluted serum for 1 hour, and 1 mL of Vero E6 medium was added after the incubation. Cells were subsequently incubated further for 10 days at 37 °C and 5% CO_2_ and monitored for cytopathic effect (CPE) by optical microscopy.

### Immunofluorescence assay (IFA) for IgG against SARS-CoV-2

Vero E6 cells were infected with SARS-CoV-2 (isolate SARS-CoV-2/human/SWE/01/2020, accession number MT093571) for 24 hours, trypsinized and mixed with uninfected Vero E6 cells, and then seeded on microscope slides. Twelve hours later, slides were fixed in acetone and stored at −80°C until further use. Serum samples were heat-inactivated at 56°C for 30 minutes prior to analysis. For analysis of total SARS-CoV-2 IgG antibody titers, serum samples were serially diluted from 1:20 to 1:5120. 25 µL of diluted serum was then added to fixed cells and incubated at 37°C for 30 min, after which the slides were washed in NaCl for 30 min. Bound SARS-CoV-2 IgG antibodies were then detected by incubating for 30 min at 37°C with a secondary AF488-conjugated AffiniPure goat anti-human IgG antibody (Jackson Immunoresearch), diluted 1:200 in 0.1% Evan’s Blue. SARS-CoV-2 IgG positive cells were visualized using a Nikon Eclipse Ni fluorescence microscope (x40 magnification). The titer of IgG in each serum sample was determined by the inverted dilution factor value for the highest dilution with positive staining.

### ELISAs

SARS-CoV-2 specific IgG and IgA antibodies in serum were detected using anti-SARS-CoV-2 ELISA kits (both from Euroimmun), according to the manufacturer’s instructions. SARS-CoV-2 specific IgM antibodies were detected using EDI Novel Coronavirus COVID-19 IgM ELISA kit (Epitope Diagnostics), according to the manufacturer’s instructions. Serum samples were heat-inactivated at 56°C for 30 minutes prior to analysis.

IL-6 levels in serum from patients and healthy controls were measured in freshly thawed serum using human IL-6 ELISA development kit (Mabtech), according to the manufacturer’s instructions. Serum samples were diluted 1:2 in ready-to-use ELISA diluent (Mabtech) prior to performing the IL-6 ELISA assay.

### Micro-neutralization assay

Two-fold dilution series from 1:10 to 1:10240 in EMEM (Gibco) + 5% FCS (Thermo Fisher Scientific) were performed on the serum samples, which were previously heat inactivated at 56°C for 30 minutes. Each dilution was subsequently mixed with equal volume of 4000 TCID_50_/ml SARS-CoV-2 (50 µl serum plus 50 µl virus) and incubated for 1 hour at 37 °C and 5% CO_2_. Each sample was prepared in duplicates. After incubation, the mixtures were added on confluent Vero E6 cells seeded on 96-well plates and incubated at 37 °C 5% CO_2_. Four days later the cells were inspected for signs of cytopathic effect (CPE) by optical microscopy. Each well was scored as either ‘neutralizing’ if less than 50% of the cell layer showed signs of CPE, or ‘non-neutralizing’ if ≥50% CPE was observed. Results are shown as the arithmetic mean of the reciprocals of the highest neutralizing dilutions from the two duplicates for each sample.

### Real-time RT-PCR

RNA was extracted from freshly thawed serum samples using the MagDEA Dx SV reagent kit and the magLEAD instrument (Precision System Science). The assay used to detect SARS-CoV-2 RNA was modified from (14): forward primer 5’-CATGTGTGGCGGTTCACTATATGT-3’, reverse primer 5’-TGTTAAARACACTATTAGCATAWGCAGT-3’, and RdRp_SARSr-P2 probe. The assay was carried out in 25 µL reaction mixtures containing 5µL RNA template, TaqMan Fast Virus 1-Step Master Mix, 0.6 µM forward primer, 0.8 µM reverse primer, and 0.2 µM probe (Applied Biosystems). Thermal cycling was performed at 50°C for 5 min, 95°C for 20 sec, followed by 40 cycles of 95°C for 3 sec, and 60°C for 30 sec in a StepOne Plus real-time PCR (Applied Biosystems).

### Statistics and data analysis

Statistical analyses were performed using GraphPad Prism software 7.0 for MacOSX (GraphPad Software). Correlation analyses were performed using Spearman’s correlation test. Statistical significance for differences between COVID-19 patients and healthy controls was determined by two-sided Mann-Whitney *U* test. P values of < 0.05 were considered statistically significant. FlowJo software version 10.5.3 (Tree Star) was used to analyze all flow cytometry data. FluoroSpot data was analyzed with Apex software (Mabtech).

## Results

Twenty COVID-19 patients were enrolled in this study during hospitalization at Karolinska University Hospital, Sweden. All twenty patients exhibited typical COVID-19 symptoms of fever, cough and breathing difficulties, and a few also experienced chest pain, myalgia and diarrhea (Table I). Some of the patients presented with common co-morbidities such as overweightness, hypertension, asthma, cardiovascular disease and diabetes mellitus type II. According to the guidelines for diagnosis and treatment of COVID-19 (Version 7) released by the National Health Commission and State Administration of Traditional Chinese Medicine, 19 out of the 20 patients were classified as severe, and 1 out of the 20 as critically ill (15). While all patients enrolled in this study were hospitalized at the infectious diseases wards at the time of sampling, two patients were treated at the intensive care unit (ICU) before inclusion in this study, while one patient required ICU treatment after the inclusion. Peripheral blood samples from the twenty COVID-19 patients were collected at a median of 15 days after the self-reported onset of symptoms, and at a median of 3 days after hospitalization (Table I, Fig. 1A and B). In parallel, seven donors without an ongoing respiratory disease or signs of inflammation were included in the study as healthy controls.

**Figure 1.**
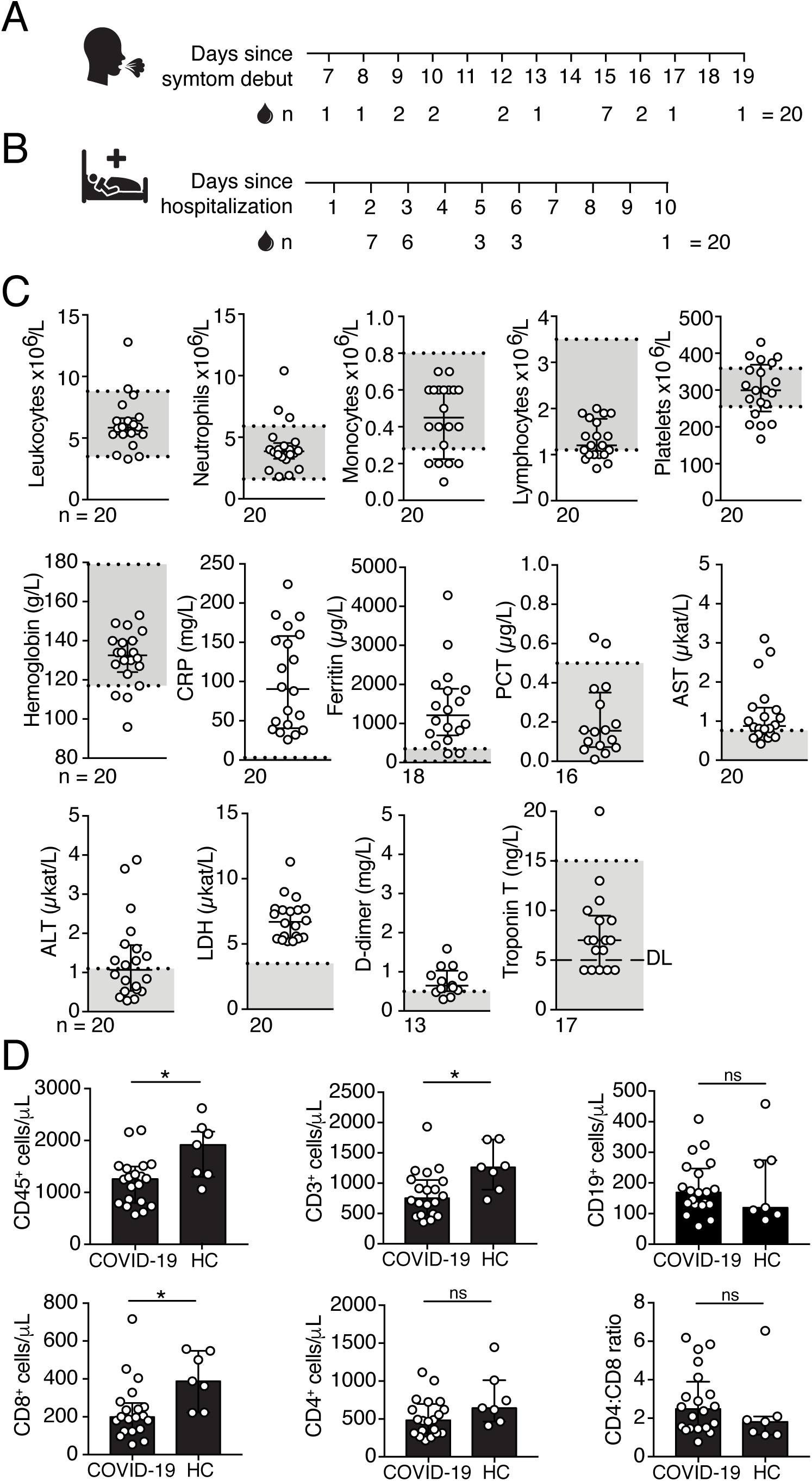
Clinical chemistry parameters in peripheral blood of COVID-19 patients. (A) Number of days since the onset of COVID-19 symptoms that peripheral blood samples were taken. (B) Number of days since hospitalization that peripheral blood samples were taken. (C) Blood cell counts and clinical chemistry parameters in peripheral blood of COVID-19 patients measured on the day of the inclusion in this study (+/– 24 hours). Gray boxes indicate the range for reference values. (D) Absolute numbers of CD45^+^, CD3^+^, CD4^+^, CD8^+^, and CD19^+^ cells in peripheral blood of COVID-19 patients (n = 20) and healthy controls (HC) (n = 7) measured by flow cytometry. Statistical significance was determined using Mann-Whitney *U* test (D). Graphs display median and IQR. n = number of patients. CRP – C-reactive protein; LDH – lactate dehydrogenase; AST – aspartate transaminase; ALT – alanine transaminase; PCT – procalcitonin; DL – detection limit. *, P < 0.05; **, P < 0.01; ***, P < 0.001; ns – not significant.

### Clinical chemistry parameters in COVID-19 patients

Blood cell counts and clinical chemistry parameters in peripheral blood of COVID-19 patients were measured at the Diagnostics Laboratory of the Karolinska University Hospital on the same day as the inclusion in this study (+/– 24 hours). Clinical chemistry tests showed increased levels of inflammatory markers, including C-reactive protein (CRP), lactate dehydrogenase (LDH) and ferritin (Fig. 1C). Some patients also presented with increased levels of aspartate transaminase (AST), alanine transaminase (ALT) and D-dimer, while hemoglobin, procalcitonin (PCT) and troponin T levels fell within the range of reference values in the majority of patients (Fig. 1C). Numbers of total leukocytes, neutrophils, monocytes and platelets in peripheral blood were within the normal range in the majority of COVID-19 patients at the time of sampling (Fig. 1C). In contrast, we observed reduced lymphocyte numbers, with the median lymphocyte numbers falling below or within the lower range of reference values (Fig. 1C). Reduced lymphocyte numbers in peripheral blood of COVID-19 patients were confirmed by flow cytometry analysis of absolute cell counts. We found significantly lower CD45^+^ lymphocyte numbers in the patients compared to healthy controls (Fig. 1D). CD3^+^ T cell and CD3^+^CD8^+^ T cell numbers were also significantly lower in patients compared to the controls, while no significant difference in the numbers of CD3^+^CD4^+^ T cells and CD19^+^ B cells was observed between COVID-19 patients and healthy controls (Fig. 1D).

### T cells and B cells are activated in COVID-19 patients

We next assessed the overall activation level of the adaptive immune system in COVID-19 patients. T cell and B cell activation levels were measured based on the expression of the activation marker CD38 and the proliferation marker Ki-67 using multicolor flow cytometry on freshly isolated peripheral blood mononuclear cells (PBMCs) (Fig. 2A). We observed significantly higher frequencies of activated CD4^+^ T cells, CD8^+^ T cells and CD19^+^ B cells in COVID-19 patients compared to healthy controls (Fig. 2B). Additionally, there was a strong positive correlation between the frequencies of activated CD8^+^ T cells and CD4^+^ T cells in COVID-19 patients (r_s_ = 0.699, P < 0.001) (Supplementary Fig. 2A).

**Figure 2.**
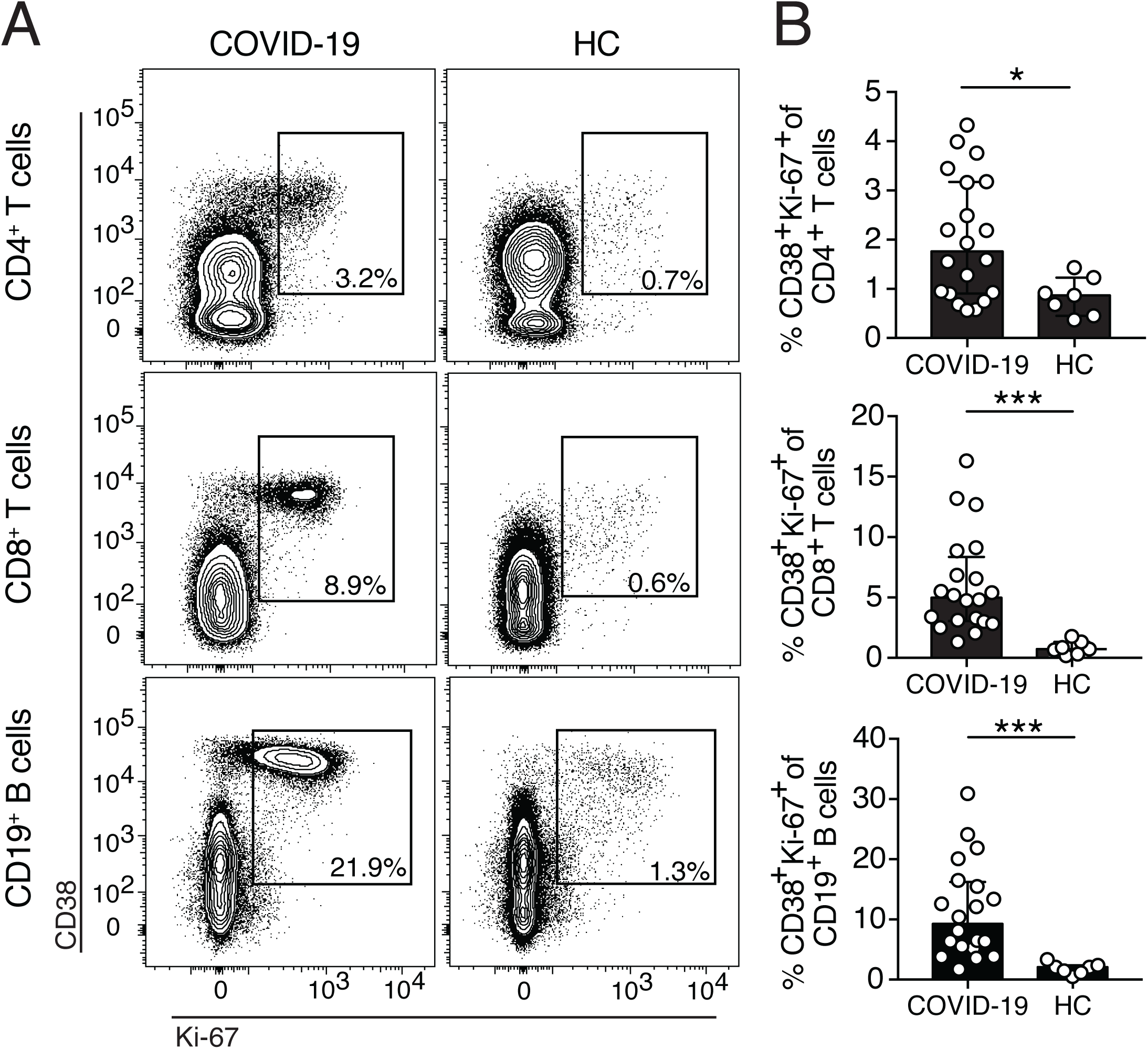
T cells and B cells are activated in peripheral blood of COVID-19 patients. (A) Representative flow cytometry plots of CD38 and Ki-67 co-expression in CD4^+^ and CD8^+^ T cells, as well as CD19^+^ B cells in one representative COVID-19 patient (16 days after symptom onset) and one healthy control (HC). (B) Frequencies of CD38 and Ki-67 co-expressing CD4^+^ T cells, CD8^+^ T cells and CD19^+^ B cells in COVID-19 patients (n = 20) and healthy donors (n = 7). Bar graphs display median and IQR. Statistical significance was determined using Mann-Whitney *U* test. *, P < 0.05; ***, P < 0.001.

### Significant expansion of antibody-secreting cells in COVID-19 patients

Flow cytometric analyses revealed increased frequencies of activated CD38^+^Ki-67^+^ B cells in the COVID-19 patients (Fig. 2A and B). Therefore, we further analyzed if antibody-secreting cell (ASC) frequencies were also increased in peripheral blood. ASCs were identified as isotype-switched B cells expressing high levels of surface CD38 and CD27 (CD19^+^CD20^low/–^ IgD^−^CD38^high^CD27^high^) (Fig. 3A). As expected, low frequencies and numbers of ASCs were detected in peripheral blood of healthy controls (Fig. 3B and C). However, significantly higher ASC frequencies and numbers were observed in COVID-19 patients compared to healthy controls, constituting up to 31% of all circulating B cells (Fig. 3B and C).

**Figure 3.**
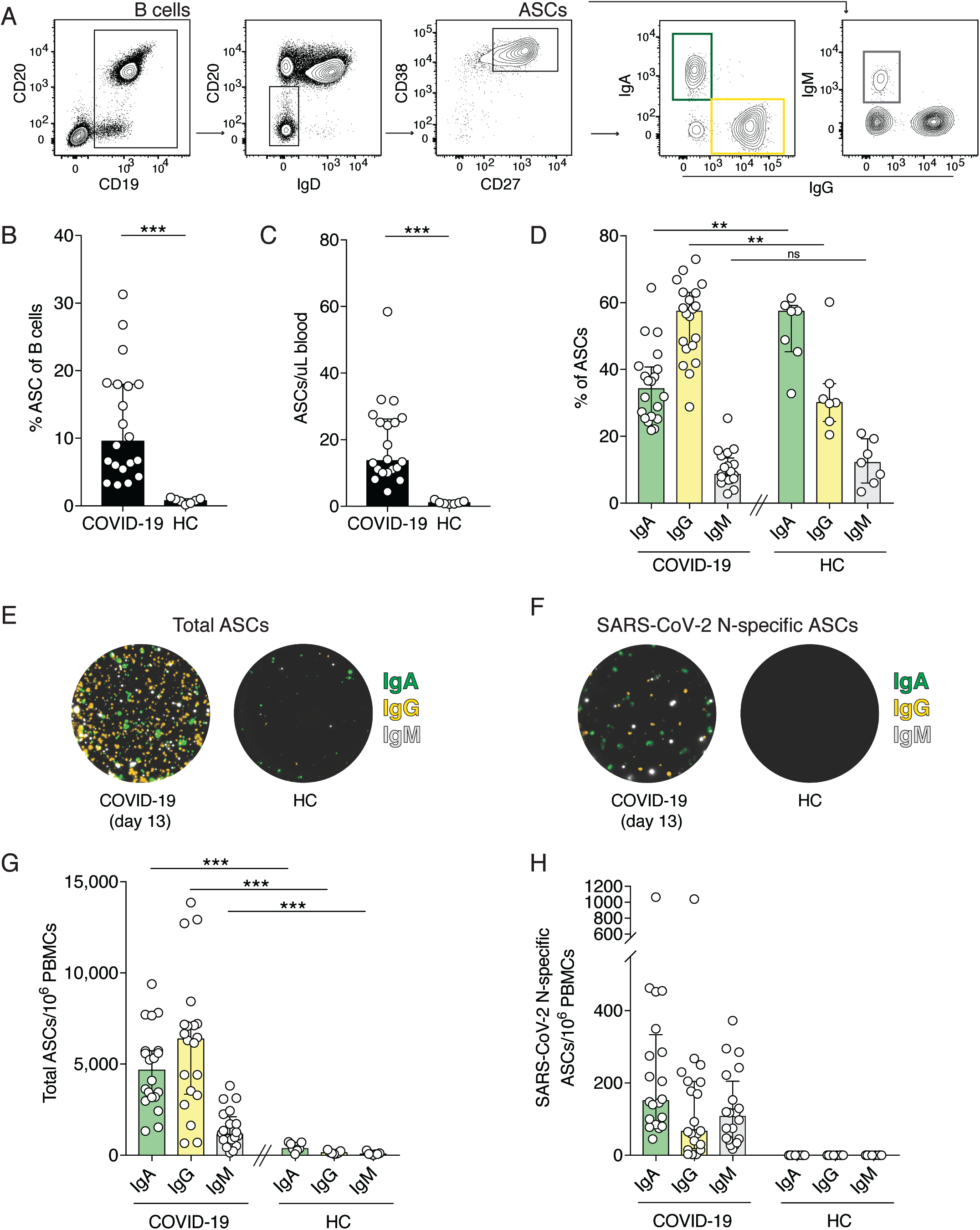
SARS-CoV-2 nucleocapsid protein-specific antibody-secreting cells (ASCs) expand in COVID-19 patients. (A) Flow cytometry gating strategy used to identify ASCs and the IgA-, IgG-, and IgM-ASC subsets. B cells are gated on live CD14^−^CD123^−^CD3^−^CD4^−^ cells. (B) Frequencies of ASCs within all B cells in COVID-19 patients and in HCs. ASCs were defined as CD19^+^CD20^low/–^IgD^−^CD38^high^CD27^high^. (C) Numbers of ASCs per microliter of whole blood, calculated using absolute B cell numbers and frequencies of ASCs measured by flow cytometry. (D) Frequencies of IgA-, IgG-, and IgM-ASCs within the total ASC population measured by flow cytometry. (E and F) Representative images of wells from a FluoroSpot assay showing total IgA-, IgG-, and IgM-ASCs (E), and SARS-CoV-2 nucleocapsid protein-specific ASCs (F) from one COVID-19 patient (13 days after symptom onset) and one healthy control. IgM fluorescence is originally red, but replaced with white in this figure for visualization purpose. (G) Numbers of total IgA-, IgG-, and IgM-ASCs per million PBMCs, as measured by FluoroSpot assay. (H) Numbers of SARS-CoV-2 nucleocapsid protein-specific IgA-, IgG-, and IgM-ASCs per million PBMCs. Experiments were performed on all COVID-19 patients (n = 20) and healthy controls (n = 7). Statistical significance was determined using Mann-Whitney *U* test (B, C, D and G). Bar graphs display median and IQR. **, P < 0.01; ***, P < 0.001; ns – not significant.

Next, we assessed the expression of immunoglobulins (Ig) by the ASCs in COVID-19 patients and healthy controls (Fig. 3A). IgA has previously been shown to be expressed by 80% of all ASCs at steady state (16). In agreement with this, the largest proportion within the ASC compartment in healthy controls was IgA^+^ ASCs, whereas the majority of ASCs in COVID-19 patients expressed IgG (Fig. 3D). IgM^+^ ASCs were present at comparable frequencies in both COVID-19 patients and healthy controls (Fig. 3D). Noteworthy, a substantial ASC expansion was detected as early as 7 days, and as late as 19 days after the onset of symptoms (Supplementary Fig. 2B).

### Detection of SARS-CoV-2-specific antibody-secreting cells in COVID-19 patients

The expansion of ASCs is usually a result of an ongoing infection and is characterized by high specificity towards the infectious agent (17). To investigate if the ASCs identified by flow cytometry were SARS-CoV-2-specific, we developed a FluoroSpot assay – a fluorescence-based variant of the enzyme-linked immunospot (ELISpot) assay, which allows for simultaneous detection of total and antigen-specific IgA-, IgG- and IgM-ASCs. To detect SARS-CoV-2-specific IgA-, IgG- and IgM-ASCs, we generated recombinant SARS-CoV-2 nucleocapsid protein (N protein).

The FluoroSpot assay confirmed the ASC expansion detected by flow cytometry, as the total numbers of IgA-, IgG-, and IgM-ASCs in COVID-19 patients were significantly higher than in healthy controls (Fig. 3E and G). Moreover, the total ASC frequencies and numbers detected by flow cytometry positively correlated with total numbers of ASCs measured by FluoroSpot (r_s_ = 0.537, P = 0.01; and r_s_ = 0.636, P = 0.003, respectively). Consistent with flow cytometry data, IgG-ASCs constituted the predominant subset of all ASCs in COVID-19 patients, followed by IgA-ASCs (Fig. 3D and G).

Importantly, we detected SARS-CoV-2 N protein-specific ASCs in all twenty COVID-19 patients, but not in controls, suggesting an active SARS-CoV-2-specific B cell response in acute COVID-19 (Fig. 3F and H). We detected IgA-, IgG- and IgM-ASCs that are specific for SARS-CoV-2 N-protein, yet at highly variable numbers among the patients. Overall, N-protein-specific ASCs constituted a small fraction of total ASCs [median (IQR) = 3.47% (2.5-4.9)]. Nonetheless, total ASC frequencies determined by flow cytometry positively correlated with N protein-specific ASC numbers (r_s_ = 0.574, P = 0.008), suggesting that the ASC expansion detected by flow cytometry may reflect the magnitude of SARS-CoV-2 N protein-specific ASC response.

### Generation of SARS-CoV-2-specific and neutralizing antibodies in COVID-19 patients

The expansion of SARS-CoV-2-specific ASCs in all of the patients in our cohort suggested that the patients had developed SARS-CoV-2-specific antibodies in response to the infection. To investigate this in detail, we next analyzed SARS-CoV-2-specific antibody levels.

First, we measured SARS-CoV-2 spike S1-specific IgA and IgG, as well as N-protein-specific IgM antibody levels by ELISAs. We found detectable SARS-CoV-2-specific IgA (15/20 patients), IgG (15/20 patients) and IgM (16/20 patients) antibody levels in most of the COVID-19 patients (Fig. 4A and B). Next, we determined total anti-SARS-CoV-2 IgG antibody levels measured towards SARS-CoV-2-infected cells using an immunofluorescence assay (IFA) (Fig. 4C). We found that 16 out of the 20 patients were positive in this assay, with titers ranging from 40 to 5120 (Fig. 4C). As expected, none of the healthy controls were positive in any of the antibody assays (Fig. 4A). Higher IgG levels were detected in patients who were sampled late compared to early after the onset of symptoms, as total and spike S1-specific SARS-CoV-2 IgG antibody levels positively correlated with the number of days since symptom onset (r_s_ = 0.577, P = 0.01, and r_s_ = 0.603, P = 0.005, respectively) (Supplementary Fig. 2C and D).

**Figure 4.**
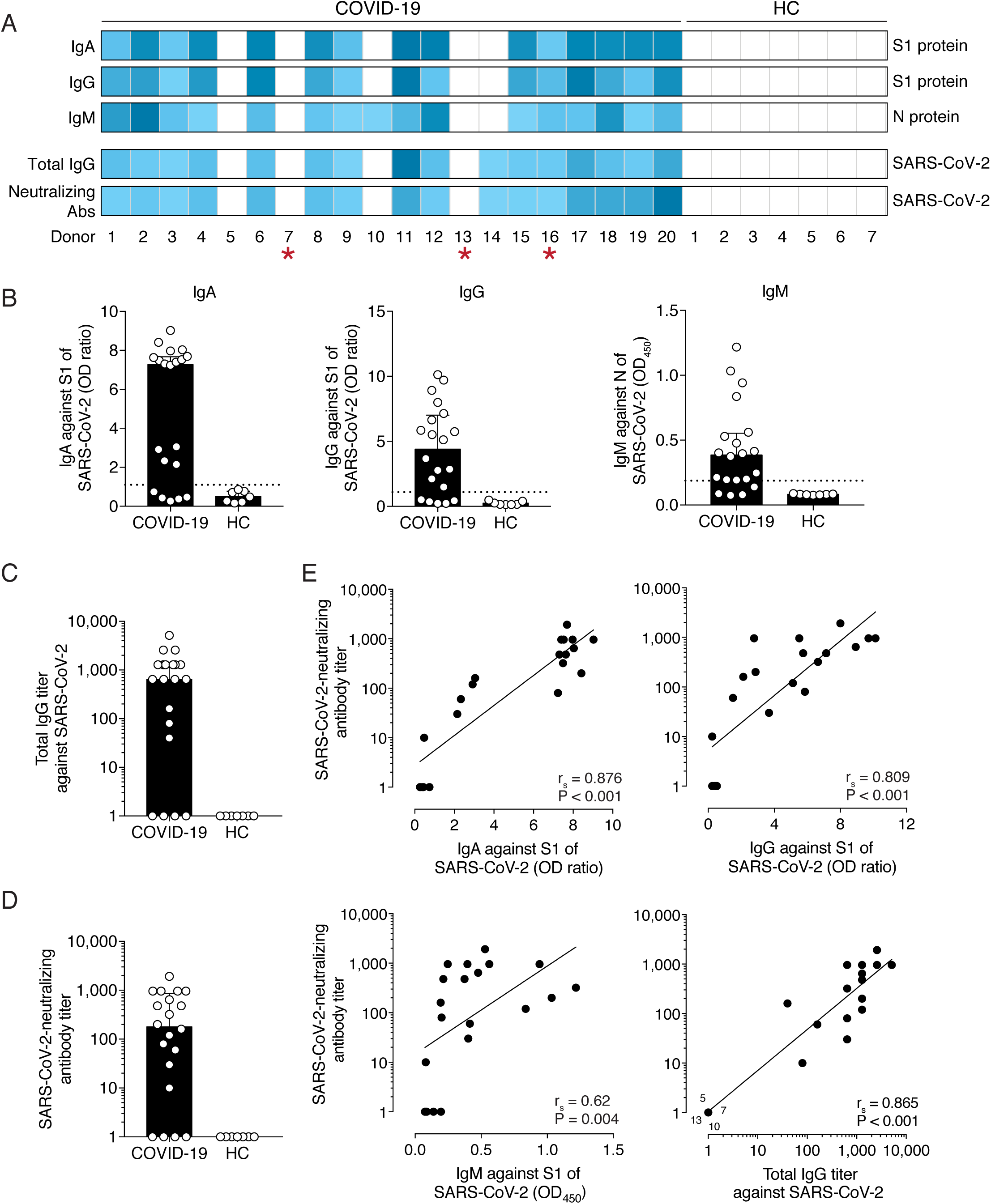
SARS-CoV-2-specific and neutralizing antibody levels in COVID-19 patients. (A) Individual antibody responses to SARS-CoV-2 in COVID-19 patients (n = 20) and healthy controls (HC) (n = 7). Positivity for IgA against S1-protein, IgG against S1-protein, IgM against N protein, total IgG antibodies against whole SARS-CoV-2, and SARS-CoV-2-neutralizing antibody titers are presented in the heatmap. Blue color indicates a positive response and the color scale is adjusted for the minimum positive assay value and the highest value recorded within the patient cohort for each assay. White boxes indicate values below positive threshold or below detection level for each assay. Red asterisks represent patients with detectable levels of SARS-CoV-2 RNA in serum. (B) IgA, IgG, and IgM antibody levels in COVID-19 patients and controls, analyzed by ELISAs. Dotted horizontal line indicates the threshold for positive result. OD, optical density. OD ratio = OD of the sample divided by OD of the calibrator. (C) Total SARS-CoV-2 IgG antibody titers determined by immunofluorescence assay. Patients with titers < 20 were assigned a value of 1. (D) SARS-CoV-2-neutralizing antibody titers determined by microneutralization assay. Patients with titers < 10 were assigned a value of 1. (E) Correlation between SARS-CoV-2-specific IgA, IgG and IgM antibody titers and SARS-CoV-2-neutralizing antibody titers, tested with Spearman’s correlation test. r_s_: Spearman’s rank correlation coefficient. P < 0.05 was considered statistically significant. Four data points from COVID-19 patients with identical values are highlighted (5, 7, 10 and 13). Bar graphs display median and IQR.

Next, using a micro-neutralization assay we assessed if the COVID-19 patients had generated SARS-CoV-2-neutralizing antibodies (18). This assay measures the ability of COVID-19 patient serum to reduce the cytopathic effect (CPE) caused by SARS-CoV-2 infection of susceptible cells *in vitro*. Neutralizing antibodies were detected in most of the patients (16/20), but at heterogeneous titers ranging from 10 to 1920 (Fig. 4D). Three of the four patients with undetectable levels of SARS-CoV-2-neutralizing antibodies were also below the level of detection for SARS-CoV-2-specific antibodies in IFA and all ELISAs (Fig. 4A).

Interestingly, we observed a strong positive correlation between total SARS-CoV-2-specific IgG antibody levels and neutralizing antibody titers in COVID-19 patients (Fig. 4E). Both S1-specific IgA and IgG, as well as N-specific IgM levels also correlated with SARS-CoV-2-neutralizing antibody titers suggesting that SARS-CoV-2-specific antibody titers may reflect the levels of neutralizing antibodies during the acute phase of COVID-19 (Fig. 4E).

Next, we assessed if the frequencies of activated T cells and B cells correlate with SARS-CoV-2-neutralizing antibody responses. We found that CD8^+^ T cell activation level, but not CD4^+^ T cell or CD19^+^ B cell activation level, positively correlated with neutralizing antibody titers (r_s_ = 0.544, P = 0.01; r_s_ = 0.271, P = 0.25, and r_s_ = −0.136, P = 0.57, respectively) (Supplementary Fig. 2E – G).

### Detection of SARS-CoV-2 RNA in serum of COVID-19 patients

An important role of neutralizing antibodies is to limit the spread of the virus. We screened all serum samples by real time RT-PCR for SARS-CoV-2 RNA (14). Three COVID-19 patients were positive for SARS-CoV-2 RNA in serum (Fig. 4A). Notably, 2 out of the 3 SARS-CoV-2 RT-PCR-positive patients lacked detectable levels of neutralizing antibodies, which might allow for a more efficient spread of SARS-CoV-2 in those patients (Fig. 4A). However, our attempts to isolate live SARS-CoV-2 from patient serum on Vero E6 cells were unsuccessful (data not shown), suggesting absence or low levels of live SARS-CoV-2 in serum of COVID-19 patients.

### Inflammatory makers, neutralizing antibodies and disease severity in COVID-19

The pro-inflammatory cytokine IL-6 and CRP serum levels have been shown to correlate with disease severity in COVID-19 patients (19, 20). We measured IL-6 serum concentration in all COVID-19 patients at the time of inclusion in this study using ELISA. We found heterogenous, yet significantly higher median concentration of IL-6 in COVID-19 patients compared to the controls (Fig. 5A).

**Figure 5.**
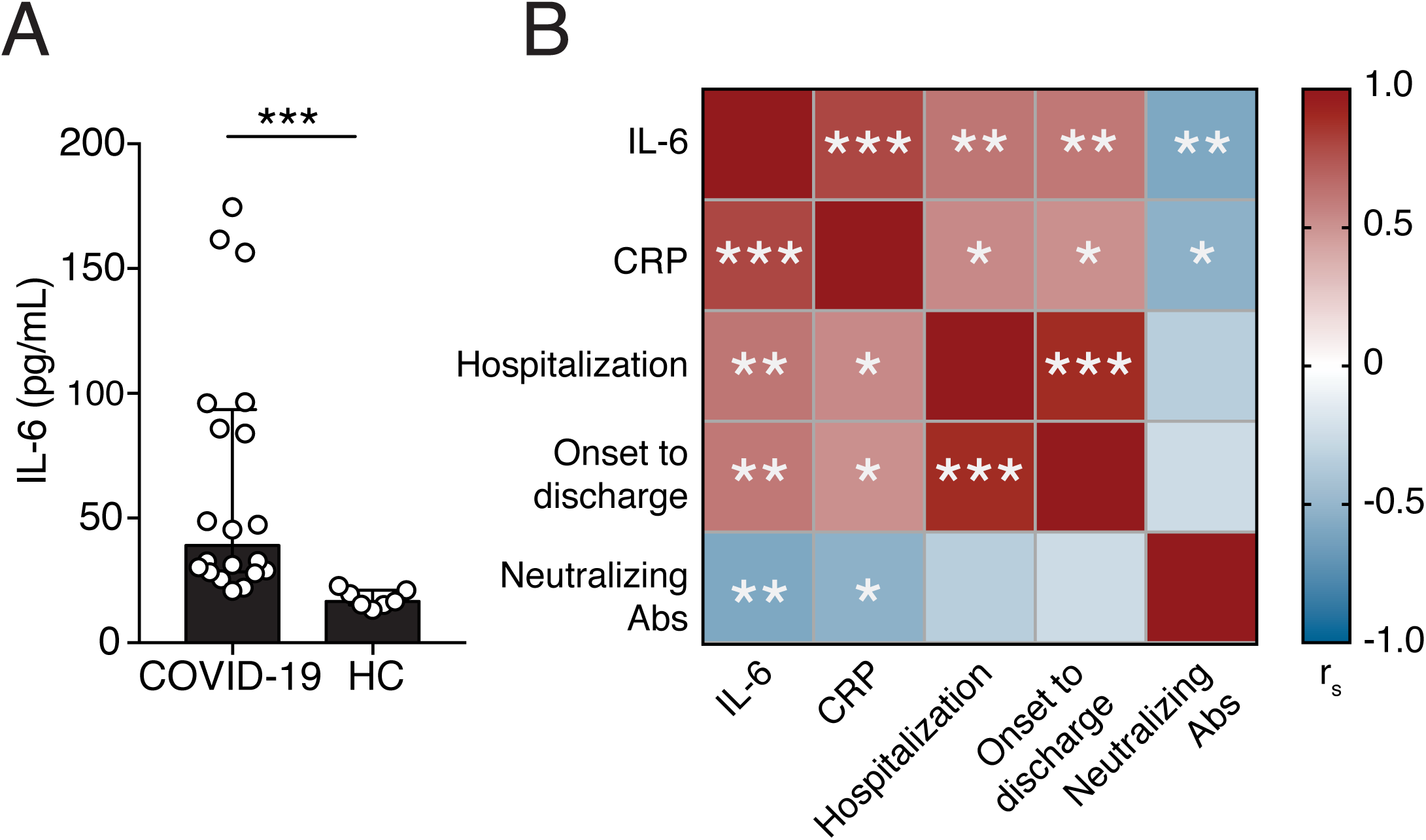
Correlations between the levels of IL-6, CRP, neutralizing antibodies and the duration of COVID-19. (A) Serum concentrations of IL-6 in COVID-19 patients and healthy controls. Bar graph displays median and IQR. Statistical significance was determined using Mann-Whitney *U* test. (B) Correlation matrix for serum IL-6, CRP, neutralizing antibody titers, duration of hospitalization and the number of days between symptom onset to discharge from hospital. Correlations were examined by Spearman’s correlation test. The color scale represents the range values for Spearman’s rank correlation coefficient (r_s_). *, P < 0.05; **, P < 0.01; ***, P < 0.001.

There was a strong positive correlation between the IL-6 and CRP levels at the time of inclusion in this study (Fig. 5B). Higher serum concentrations of both IL-6 and CRP were higher in patients with longer duration of hospitalization (Fig. 5B). Notably, IL-6 and CRP serum concentrations negatively correlated with neutralizing antibody titers in COVID-19 patients, indicating a possible link between inflammation and humoral responses in COVID-19 (Fig. 5B).

## Discussion

Here, we demonstrated that COVID-19 patients elicit a SARS-CoV-2-specific B cell response, indicated by the expansion of SARS-CoV-2-specific ASCs. Although not all patients in the present cohort had detectable levels of SARS-CoV-2-specific antibodies at the time of sampling, SARS-CoV-2 nucleocapsid protein-specific ASCs could be detected in all patients using the FluoroSpot assay. In addition, we showed a clear relationship between the levels of SARS-CoV-2-specific antibodies and total SARS-CoV-2-neutralizing antibodies. This suggests that standard serological assays may inform about the levels of neutralizing SARS-CoV-2-specific antibodies, which may offer protection from a re-infection. Additionally, tools employed in this study may be of relevance in the assessment of long-lasting immunity after SARS-CoV-2 infection and vaccination.

The expansion of ASCs in the present cohort of COVID-19 patients highlights an active B cell response to SARS-CoV-2 infection. The frequencies of ASCs within the total circulating B cell population were highly variable between COVID-19 patients, ranging from 3 to 31%. Although the patients in this cohort were sampled at different timepoints following the onset of symptoms, a high variability in ASC frequencies have also been observed among patients infected with influenza or dengue virus sampled at normalized timepoints (e.g., 7 days after febrile illness) (4, 5). Although ASCs have been shown to expand to very high levels in some viral infections, constituting up to 50-80% of all peripheral B cells, the highest ASCs frequency in this cohort of COVID-19 patients was 31% (5, 7, 21). Healthy controls in this study had very low frequencies of ASCs in the circulation, a finding consistent with previous observations (5, 6, 16).

The ASC response in peripheral blood is usually transient and peak at around 7 days after the onset of febrile illness, dropping to steady-state levels 2-3 weeks later (8). Interestingly, we observed a substantial ASC expansion as late as 19 days following symptom debut. This finding highlights the similarity between COVID-19 and respiratory syncytial virus (RSV) infection, where a significant ASC expansion could be detected as late as 22-45 days after the onset of symptoms (9). The prolonged ASC expansion in RSV infection was associated with longer RSV shedding in the airways. Thus, the kinetics of ASC response may be dependent on pathogen persistence. In support of our findings, a recent report on longitudinal immune responses in a single COVID-19 patient with mild disease showed the peak of ASC expansion at day 8 after symptom onset, but a substantial ASCs response could still be detected by day 20 (22). Thus, it remains to be investigated whether SARS-CoV-2 persistence in the airways may lead to a prolonged ASC response in circulation.

The expansion of ASCs is usually characterized by high specificity towards the infectious agent (17). Here, we successfully demonstrated that COVID-19 patients generate SARS-CoV-2-specific ASC responses using a FluoroSpot assay. We were able to successfully identify SARS-CoV-2-specific IgA-, IgG- and IgM-ASCs in all COVID-19 patients. The frequency of SARS-CoV-2-specific ASCs within the total ASC population in this patient cohort was relatively low. However, since only the SARS-CoV-2 N protein was used to detect specific ASCs in this study, the numbers of N protein-specific ASCs do not reflect the overall magnitude of the specific ASC response. The use of SARS-CoV-2 viral lysate or recombinant SARS-CoV-2 proteins in combination would allow for the evaluation of the overall ASC response. Furthermore, the FluoroSpot assay described in this study could be adapted to evaluate memory B cell responses to various SARS-CoV-2 proteins in convalescent COVID-19 patient samples.

The protective level of SARS-CoV-2-neutralizing antibodies in COVID-19 has not yet been determined. Monoclonal antibodies isolated from convalescent COVID-19 patients showed a strong SARS-CoV-2-neutralizing capacity *in vitro* and *in vivo* in transgenic mice expressing human SARS-CoV-2 receptor ACE2 (23). However, extensive longitudinal studies in humans are required to investigate if recovered COVID-19 patients are protected from re-infection with SARS-CoV-2. Seroconversion in COVID-19 patients, measured by detectable SARS-CoV-2-specific IgG levels, has been recently shown to take place within nineteen days after the onset of symptoms (24). The four patients in the present COVID-19 cohort without detectable neutralizing antibody titers were sampled at days 7, 8, 9, and 15 after symptom debut. Although these four patients did not have detectable neutralizing antibody titers, SARS-CoV-2 specific ASCs were detected. This indicates that the FluoroSpot assay may be a more sensitive method to detect early B cell responses to SARS-CoV-2, compared to standard serological assays.

In this cohort of patients, 2 out of the 4 patients with undetectable neutralizing antibody levels had SARS-CoV-2 RNA in serum. Currently, the extent to which neutralizing antibodies contribute to the control of viremia in COVID-19 is unclear. Our results seem to suggest that neutralizing antibodies may not be the only player in SARS-CoV-2 control in circulation, as one patient with substantial levels of SARS-CoV-2-neutralizing antibodies in this cohort had detectable serum levels of SARS-CoV-2 RNA. A pilot study assessing the effectiveness of COVID-19 patient treatment with convalescent plasma containing SARS-CoV-2-neutralizing antibodies found that the treatment reduced viral load in patients within a few days after administration, suggesting that neutralizing antibodies have a role in the management of viremia in COVID-19 (25).

Rapid serology tests are essential for COVID-19 diagnostics and for national serosurveillance programs. Neutralizing antibody tests, however, are usually laborious and time-consuming. Therefore, diagnostic tests often rely on other methods such as ELISA. Antibody levels detected by ELISA, however, do not necessarily reflect the neutralization capacity of patient serum. Here, we demonstrated a strong correlation between SARS-CoV-2-specific and SARS-CoV-2-neutralizing antibody titers. These data suggest that the levels of SARS-CoV-2-specific antibodies measured with ELISA may reflect the capacity of serum from COVID-19 patients to neutralize SARS-CoV-2.

COVID-19 patients generally present with decreased lymphocyte numbers in peripheral blood, which is particularly pronounced in severe cases (19, 26, 27). T cells appear to be the most affected lymphocyte subset. In agreement with these observations, we found decreased numbers of lymphocytes, and significantly lower T cell numbers in the present cohort of COVID-19 patients.

In addition to lymphopenia, COVID-19 patients typically display increased concentrations of inflammatory cytokines in serum, especially in severe cases (19, 27). The levels of inflammatory markers IL-6 and CRP, in particular, have been shown to correlate with disease severity in COVID-19 patients (19, 20). In this cohort of COVID-19 patients we also observed increased levels of IL-6 and CRP, with comparable median values to those measured by Liu *et al*. in severe COVID-19 patients (20). We show that higher IL-6 and CRP levels were observed in patients who were hospitalized for longer, suggesting that these markers may be of use in predicting the duration of COVID-19.

Lastly, we observed an inverse correlation between IL-6 levels and SARS-CoV-2-neutralizing antibody titers in COVID-19 patients. This relationship may indicate that the lack of SARS-CoV-2-neutralizing antibodies leads to higher inflammation, or that high levels of inflammation impair or delay antibody responses. Further studies where a longitudinal patient sampling at controlled timepoints is performed are required to describe the relationship between inflammation and humoral responses in COVID-19.

Taken together, we provided a detailed description of clinical and immunological parameters in twenty COVID-19 patients, with a focus on B cell and antibody responses. We demonstrated that COVID-19 patients elicit a SARS-CoV-2-specific B cell response, indicated by the expansion of SARS-CoV-2-specific ASCs. In addition, we showed a clear relationship between the levels of SARS-CoV-2-specific antibodies and total SARS-CoV-2-neutralizing antibodies. The tools employed in this study may be of relevance in the assessment of long-lasting immunity after SARS-CoV-2 infection and vaccination.

## Acknowledgments

We would like to thank the patients and healthy controls for participating in the study. We thank the nurses at the Karolinska University Hospital, Stockholm, Sweden for blood sampling of patients, T. Aktas and M. Olausson at the Public Health Agency of Sweden for technical assistance, and P. Schierloh at IBB-UNER-CONICET (Argentina) for providing input on statistical analysis.

## 2. Abbreviations

ASC: antibody-secreting
cell CPE: cytopathic effect
COVID-19: coronavirus disease 2019
N protein: nucleocapsid protein
SARS-CoV-2: severe acute respiratory syndrome coronavirus 2
S1 protein: subunit 1 of SARS-CoV-2 spike protein

---

**Supplementary Figure 1.**
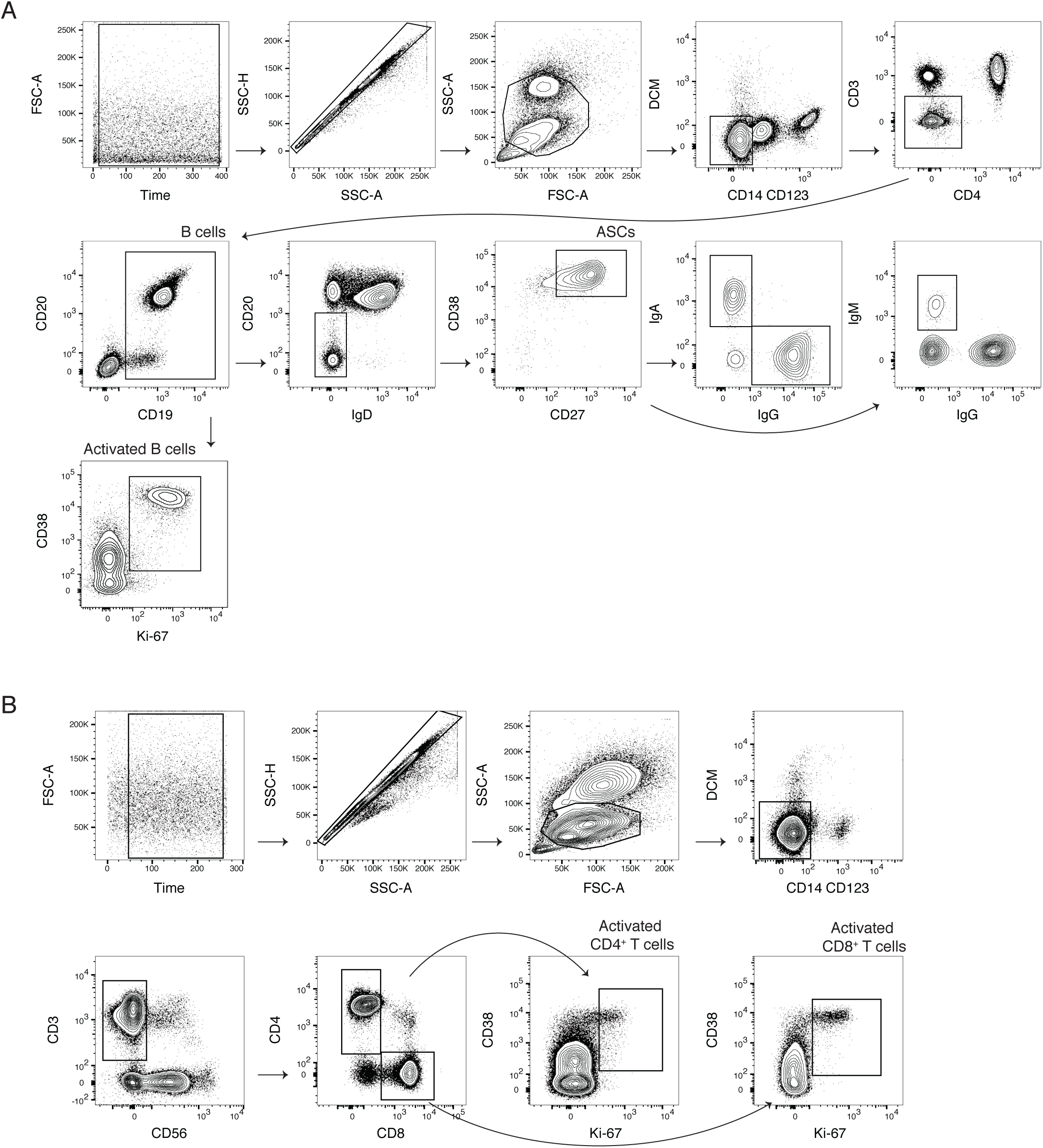
(A) Flow cytometry gating strategy for antibody-secreting cells (ASCs), the IgA, IgG and IgM ASC subsets, as well as activated B cells. (B) Flow cytometry gating strategy for CD4^+^ and CD8^+^ T cell activation, defined as the co-expression of CD38 and Ki-67.

**Supplementary Figure 2.**
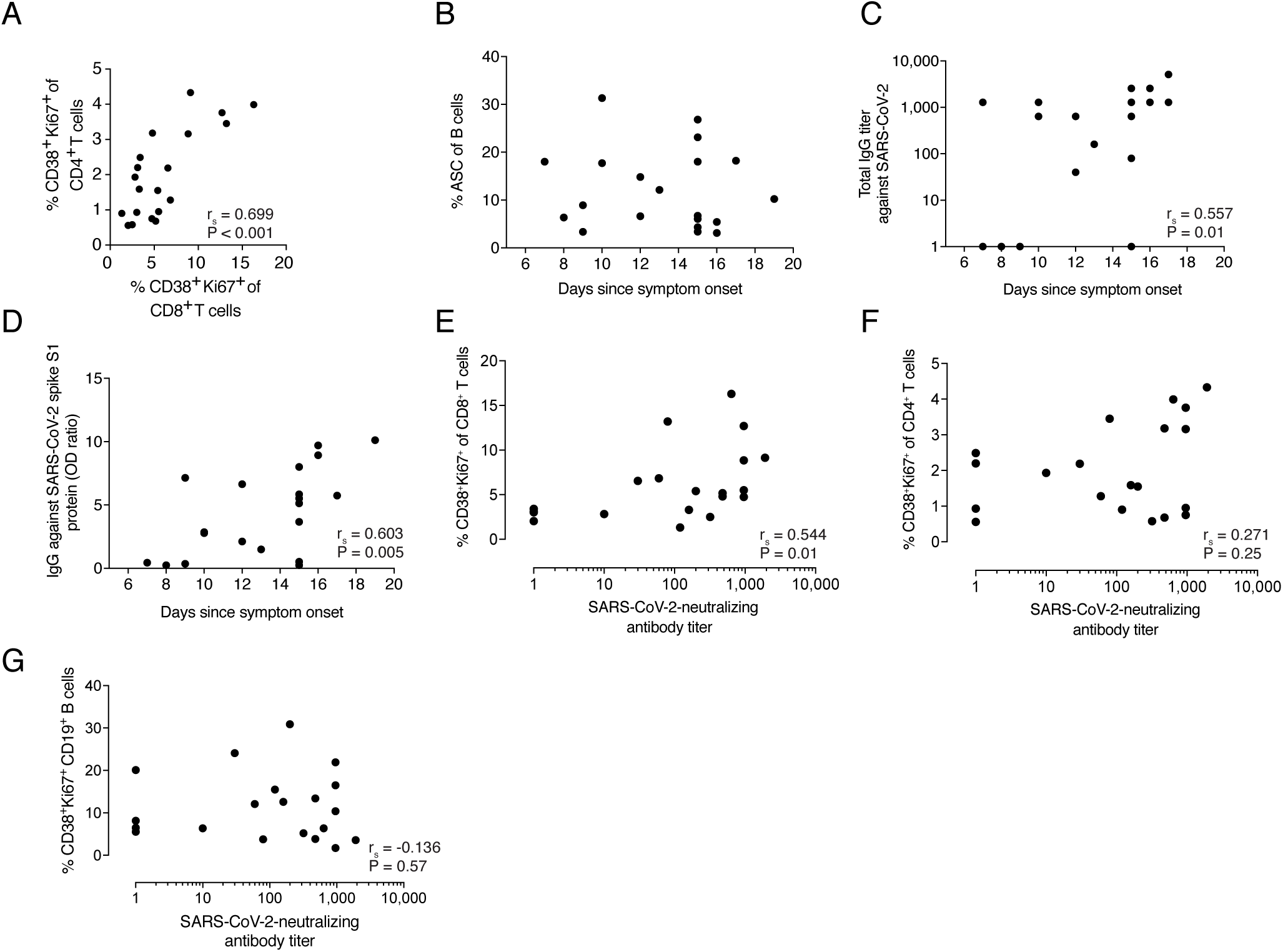
(A) Correlation between CD4^+^ and CD8^+^ T cell activation levels, as measured by CD38 and Ki-67 co-expression using flow cytometry. (B) Graph displaying the antibody-secreting cell (ASC) frequencies within the total pool of CD19^+^ B cells and the number of days since symptom onset of COVID-19. (C) Correlation between total IgG titers against SARS-CoV-2 (measured by IFA) and the number of days since symptom onset of COVID-19. (D) Correlation between levels of SARS-CoV-2 S1 protein-specific IgG levels and the number of days since symptom onset of COVID-19. (E-G) Correlations between activated T cells and B cells with SARS-CoV-2 neutralizing antibody titers. Correlations were examined by Spearman’s correlation test. r_s_: Spearman’s rank correlation coefficient. P < 0.05 was considered statistically significant.

